# Federated Learning for multi-omics: a performance evaluation in Parkinson’s disease

**DOI:** 10.1101/2023.10.04.560604

**Authors:** Benjamin Danek, Mary B. Makarious, Anant Dadu, Dan Vitale, Paul Suhwan Lee, Mike A Nalls, Jimeng Sun, Faraz Faghri

**Author notes:** Corresponding author: Dr. Faraz Faghri (FF).

## Abstract

While machine learning (ML) research has recently grown more in popularity, its application in the omics domain is constrained by access to sufficiently large, high-quality datasets needed to train ML models. Federated Learning (FL) represents an opportunity to enable collaborative curation of such datasets among participating institutions. We compare the simulated performance of several models trained using FL against classically trained ML models on the task of multi-omics Parkinson’s Disease prediction. We find that FL model performance tracks centrally trained ML models, where the most performant FL model achieves an AUC-PR of 0.876 ± 0.009, 0.014 ± 0.003 less than its centrally trained variation. We also determine that the dispersion of samples within a federation plays a meaningful role in model performance. Our study implements several open source FL frameworks and aims to highlight some of the challenges and opportunities when applying these collaborative methods in multi-omics studies.

**The Bigger Picture:** The wide-scale application of artificial intelligence and computationally intensive analytical approaches in the biomedical and clinical domain is largely restricted by access to sufficient training data. This data scarcity exists due to the isolated nature of biomedical and clinical institutions, mandated by patient privacy policies in the health system or government legislation. Federated Learning (FL), a machine learning approach that facilitates collaborative model training is a promising strategy to address these restrictions. Therefore, understanding the limitations of cooperatively trained FL models, and their performance differences to similar, centrally trained models, is crucial to valuing their implementation in the broader biomedical research community.

## Introduction

In recent years, Machine Learning (ML) algorithms have gained popularity as a possible vehicle for solving many long-standing research questions in the clinical and biomedical setting. Concretely, the adoption of sophisticated ML models can aid in tasks such as biomarker detection, disease subtyping ^1^, disease identification ^2^; ^3^; ^4^, and the development of novel medical interventions. The nascence of powerful predictive methods such as ML has enabled investigations into advanced analytics of granular patient features, such as genomics and transcriptomics ^5–7^, to achieve the ultimate goal of precision medicine.

Adopting ML methods is constrained by access to high-quality datasets. In biomedical studies, curating such quality datasets is particularly difficult due to the sample collection and processing costs and the barriers associated with recruiting patients who meet a study’s inclusion/exclusion criteria. The challenge of this task is exacerbated by the fact that institutions that typically collect biomedical samples cannot easily share human specimens’ data due to data privacy regulations, such as HIPAA, EU GDPR, India’s PDPA, Canada’s PIPEDA, mandated by medical systems, or at a national, and international level ^8^.

Federated Learning (FL) is an optimization method for performing machine learning model training among a group of clients, allowing each client to maintain governance of their local data. Initially developed for learning user behavior patterns on personal mobile devices without breaching individual privacy ^9^, FL has found valuable applications in numerous domains, including finance ^10^, medicine ^11, 12^, and the pharmaceutical industry ^13^. In biomedical research, FL represents an opportunity to enable cross-silo analytics and more productive collaboration ^14–16^.

This work evaluates Federated Learning methods’ practical availability and utility to enable large-scale, multi-institutional, and private analytics of multiple modality biomedical samples. Specifically, we aim to identify the frameworks biomedical researchers can use to perform federated learning in their work, the expected performance changes, and the implementation challenges they may face. We also discuss the opportunities and limitations of applying federated learning in multi-omics, where samples capture a patient’s genomic, transcriptomic features, and clinical and demographic information through the case study of Parkinson’s disease prediction.

We use the multi-modal Parkinson’s disease prediction as a case study for testing Federated Learning on omics data. Timely and accurate diagnosis of neurodegenerative diseases like Parkinson’s is crucial in exploring the efficacy of novel therapies to treat and manage the disease. Since the onset of these diseases typically begins many years before any visible symptoms, early detection is difficult to achieve at a clinical level alone. Usually, it requires information inherent to the patient’s biology. ^6^ have already determined that leveraging genomic and transcriptomic information as part of the diagnosis process allows for higher model performance. This study demonstrates the feasibility and practicality of deploying FL for Parkinson’s disease prediction.

## Results

### Federated learning models trained using publicly available and accessible frameworks results follow central model performance

We aim to evaluate the performance differences between classical ML algorithms frequently used in the biomedical setting, against federated learning algorithms suitable for cross-silo modeling. An overview of our approach (**Fig. 1**) shows the experimental design for evaluating several FL and central ML methods in the task of Parkinson’s disease prediction based on genomic, transcriptomic, and clinico-demographic features (**Supplementary Table 1**). The datasets used in experiments originate from studies providing clinical, demographic, and biological information of Parkinson’s Disease patients, the Parkinson’s Progression Marker Initiative (PPMI) and Parkinson’s Disease Biomarkers Program (PDBP). The PPMI dataset is a longitudinal, observational study where patients contribute clinical, demographic, imaging data, as well as biological samples for whole-genome sequencing and whole-blood RNA sequencing. PPMI specifically includes newly diagnosed and drug-naïve patients, collected at clinical sites globally over a span of 5 to 13 years. The PDBP dataset provides clinical, genetic, imaging, and biomarker data associated with Parkinson’s disease, Lewy Body Dementia, and other parkinsonisms. Patients in PDBP are not necessarily newly diagnosed or drug-naïve. The PPMI dataset is used for model training, validation, and testing. The PDBP dataset is used strictly as an external test set. In our experiments, the PPMI dataset is split into *K* folds, one of which is used as a holdout test set, and the remaining folds are used for model training and validation. To establish a baseline performance of classically trained, central algorithms representative of methods used in the current biomedical research paradigm, several central ML algorithms are fit to the training set (**Supplementary Table 2**) and tested on the holdout PPMI fold, as well as the whole PDBP dataset. To simulate the cross-silo federated setting, the training set is split into *N* disjoint subsets, referred to as client datasets. Where each baseline ML algorithm is fit to the full centralized training dataset, the FL model is fit to *N* disjoint, siloed client datasets, the union of which equates to the entire training data set. An illustration of the FL training process is shown in **Fig. 2**. The fitted FL models are finally evaluated against the PPMI test fold, and the PDBP dataset.

**Figure 1:**
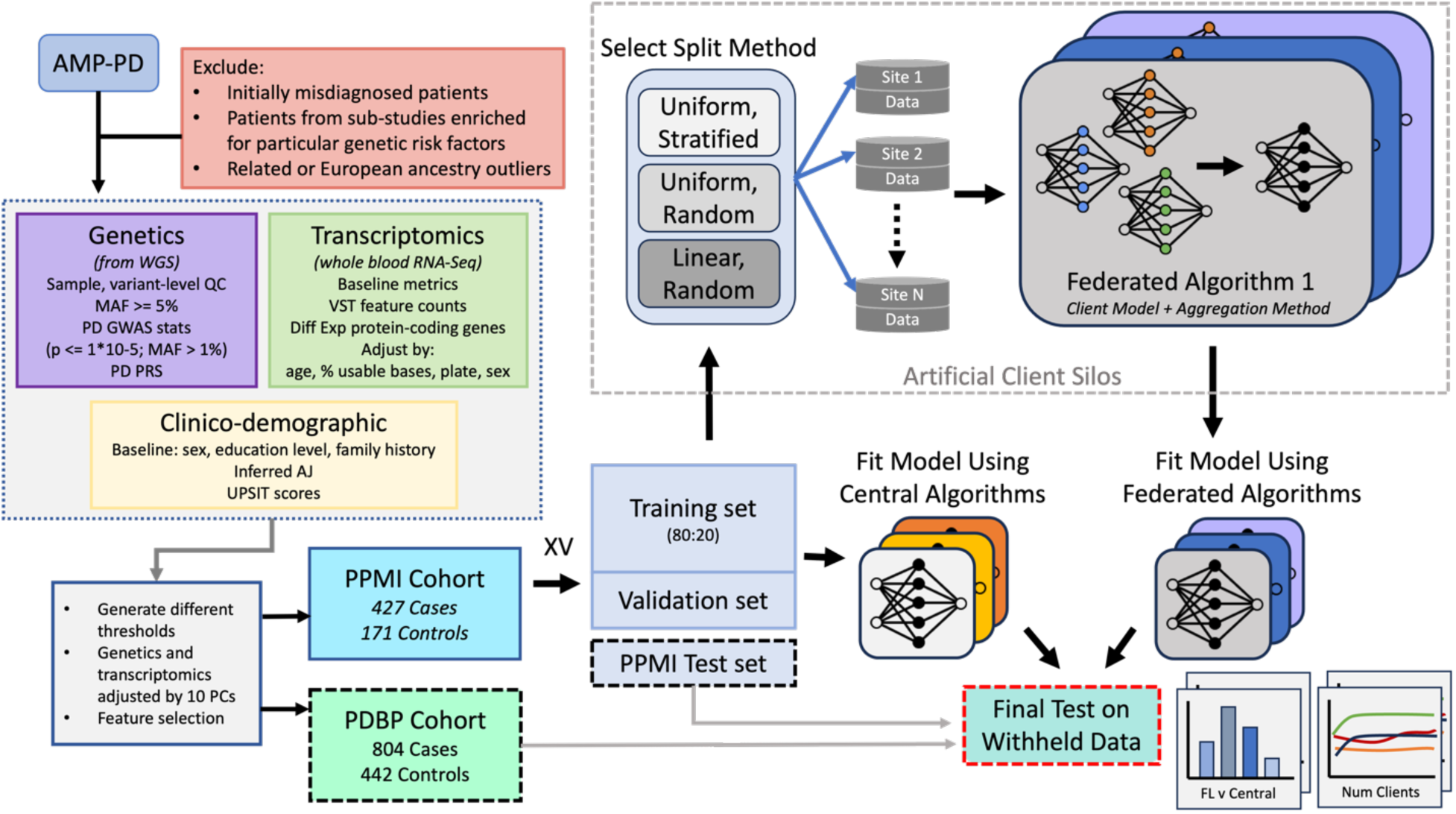
The experiment workflow diagram and the data summary. The harmonized, and joint-called PPMI and PDBP cohorts originate from the AMP-PD initiative. The PPMI cohort is split into K folds, where one fold is left as a holdout (internal) test set, and the remaining are used for model fitting. The training folds are split using an 80:20 ratio to form the training validation split. The training split is distributed among N clients using one of the split strategies to simulate the cross-silo collaborative training setting. FL Methods consist of a local learner and an aggregation method. Similarly, several central algorithms are used to fit the training data. The resultant Global FL models and the ML models resulting from central training are tested on the PPMI holdout fold (internal test) and the whole PDBP test set (external test).

**Figure 2:**
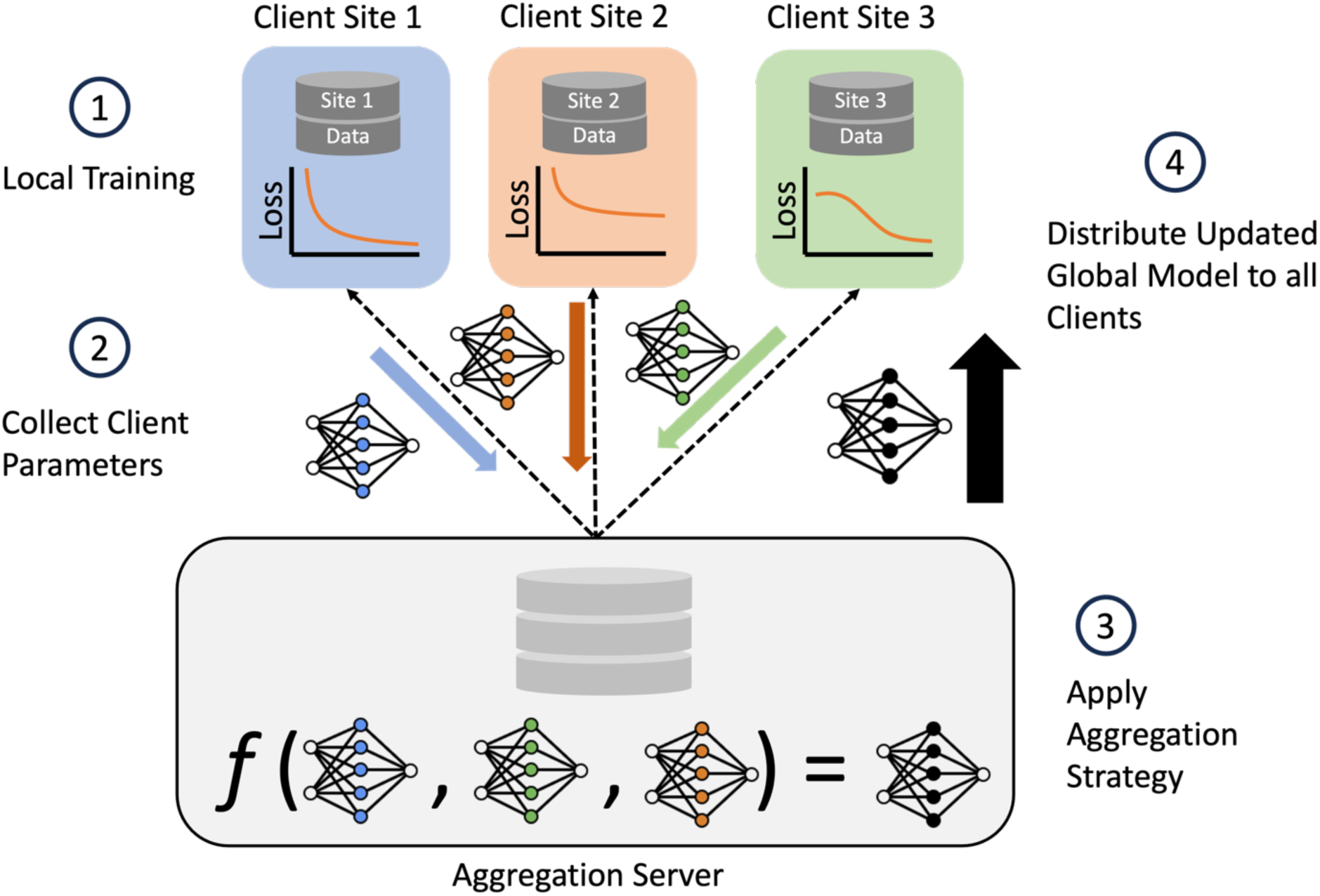
Federated architecture and training summary. The FL architecture used in the study also illustrates 1 round of FL training for the case of N=3 clients. The aggregation server aggregates trained local learner parameters from clients and computing a global model. Client Sites contain their own siloed dataset, each with different samples. The trained client parameters are represented by the blue, orange, and green weights; the black weights represent the aggregated global model. Client model aggregation implemented by the federated learning strategy is denoted by f. Once global weights are computed, a copy is sent to each client; the global model is used to initialize the local learner model weights in subsequent FL training rounds.

In the optimistic FL setting where we compare a federation of *N=2* client sites, which have been assigned samples through uniform stratified random sampling without replacement, against the central baseline algorithms (**Fig. 3**), it can be seen that for all the included FL methods, the absolute difference in performance is relatively small. For the internal test set, the central Logistic Regression (LR) Classifier ^17^ has an Area Under the Precision-Recall Curve (AUC-PR) of 0.915±0.039 (standard deviation across K = 6 folds). Among the classifiers trained using FL, which implement the same local learner, FedAvg ^18^ LR, FedProx ^19^ μ = 0.5 LR, and FedProx μ = 2 LR, have an AUC-PR of 0.874±0.042, 0.887±0.041, 0.906±0.04. In the external test set, a similar relationship between central LR and federated LR is exhibited, where classical LR has an AUC-PR of 0.842±0.009 and FedAvgLR, μ = 0.5 LR, and FedProx μ = 2 LR, have an AUC-PR of 0.826±0.011, 0.823±0.015, and 0.835±0.007. This general reduction in performance between the centrally trained classifier and the classifier trained collaboratively through FL is a trend observed in nearly all of the FL algorithms across both test sets. Similarly, the central MLP classifier ^20^ has an AUC-PR of 0.892±0.032 and 0.826±0.007 respectively, while the best performing FL variation, FedProx MLP μ = 2 has an AUC-PR of 0.868±0.06 and 0.785±0.015 for the internal and external test set respectively. A similar trend can be observed for FedAvg XGBRF ^21^, where the performance reduction is proportional to that observed in other learners. The only exception to this pattern is in the case of SGDClassifier ^22, 23^, where in the PDBP exhibits a marginal improvement of 0.002 AUC-PR. The resulting performance details, including several additional centrally trained ML algorithms, are in **Table 1**, and **Table 2**. The statistical significance of pairwise observed differences in performance is presented in **Supplementary Table 3.** The results of this side-by-side comparison of FL methods in an idealistic setting with minimal heterogeneity among client datasets show a relatively small difference in performance compared to the central algorithms.

**Figure 3:**
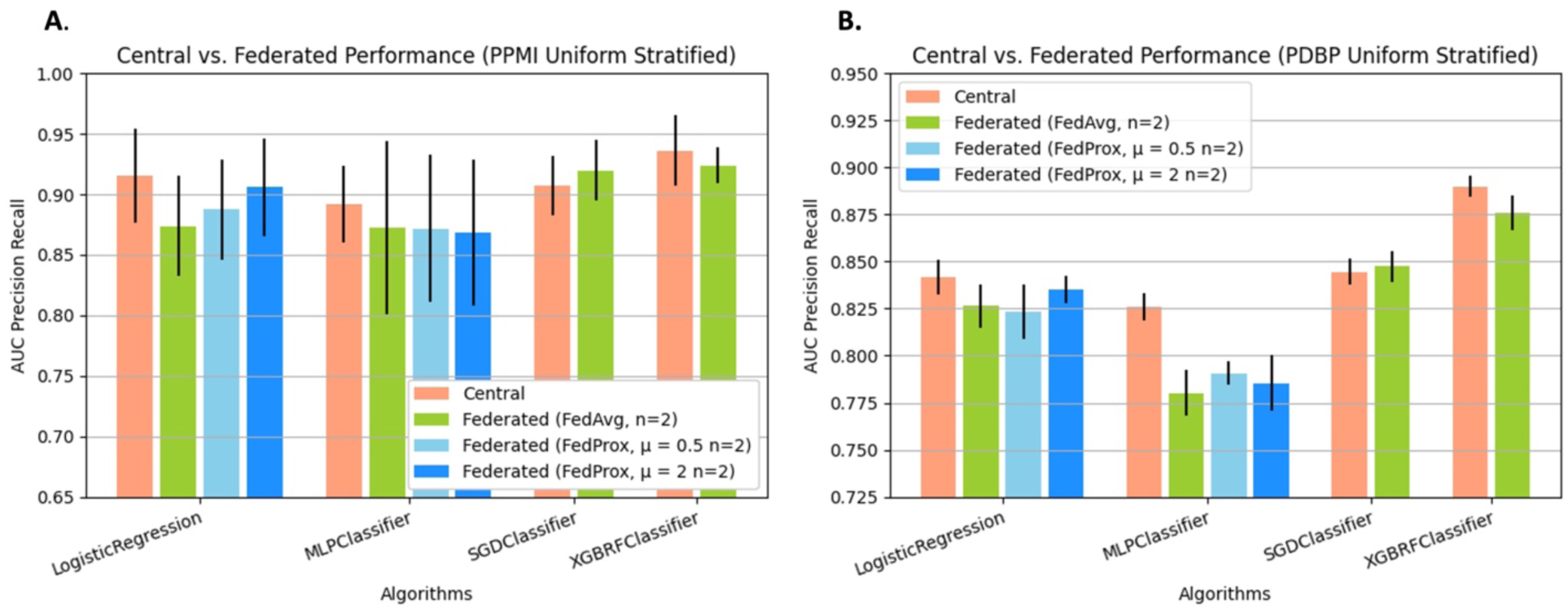
Federated learning models trained using publicly available and accessible frameworks results follow central model performance. The Precision Recall Area Under Curve (PRAUC) comparing Central Algorithms against Federated Algorithms. We pair FL algorithms with central algorithms by the local learning algorithm applied at client sites. Federated Algorithms receive the training data set split across *n=2* clients, using label stratified random sampling. Presented data is mean score and standard deviation resulting from cross validation.

**Table 1:** Performance of Several models trained using classical machine learning methods, and federated learning methods, where the number of participating clients in the federation is *N=2*, tested on the PPMI dataset. Data reported is mean and standard deviation across *K=6* fold cross validation.

| Algorithm Name | ROC-AUC | AUC-PR | Balanced Accuracy | Precision | Recall | F 0.5 | F 1 | F 2 | Log Loss | Matthews Correlation Coefficient |
| --- | --- | --- | --- | --- | --- | --- | --- | --- | --- | --- |
| AdaBoost Classifier | 0.865±0.033 | 0.939±0.01 | 0.736±0.069 | 0.844±0.043 | 0.89±0.047 | 0.897±0.017 | 0.885±0.027 | 0.942±0.013 | 0.632±0.007 | 0.499±0.122 |
| Bagging Classifier | 0.82±0.05 | 0.916±0.025 | 0.69±0.052 | 0.813±0.027 | 0.897±0.061 | 0.865±0.021 | 0.862±0.024 | 0.933±0.01 | 1.001±0.382 | 0.428±0.123 |
| GradientBoosting Classifier | 0.879±0.046 | 0.943±0.026 | 0.723±0.072 | 0.833±0.042 | 0.911±0.021 | **0.916±0.03** | 0.894±0.022 | **0.942±0.015** | 0.444±0.099 | 0.486±0.119 |
| KNeighbors Classifier | 0.61±0.099 | 0.806±0.065 | 0.533±0.029 | 0.729±0.014 | 0.937±0.046 | 0.782±0.023 | 0.837±0.007 | 0.927±0.004 | 2.836±0.617 | 0.111±0.104 |
| LinearDiscriminantAnalysis Classifier | 0.763±0.045 | 0.883±0.031 | 0.681±0.053 | 0.826±0.04 | 0.77±0.05 | 0.7±0.344 | 0.714±0.35 | 0.776±0.38 | 1.608±0.488 | 0.347±0.095 |
| LogisticRegression Classifier | 0.831±0.068 | 0.915±0.039 | 0.734±0.072 | 0.841±0.043 | 0.894±0.028 | 0.872±0.047 | 0.883±0.033 | 0.939±0.011 | 0.648±0.203 | 0.493±0.134 |
| MLP Classifier | 0.739±0.078 | 0.892±0.032 | 0.703±0.059 | 0.833±0.038 | 0.815±0.054 | 0.843±0.038 | 0.858±0.034 | 0.932±0.013 | 6.616±1.844 | 0.402±0.119 |
| QuadraticDiscriminantAnalysis Classifier | 0.504±0.057 | 0.774±0.029 | 0.504±0.057 | 0.725±0.055 | 0.385±0.081 | 0.757±0.008 | 0.833±0.006 | 0.926±0.003 | 19.674±1.492 | 0.009±0.105 |
| RandomForest | 0.816±0.076 | 0.917±0.027 | 0.552±0.034 | 0.736±0.016 | **0.993±0.017** | 0.857±0.043 | 0.874±0.032 | 0.942±0.014 | 0.508±0.029 | 0.249±0.121 |
| SGD Classifier | 0.755±0.065 | 0.907±0.025 | 0.735±0.062 | 0.846±0.032 | 0.857±0.068 | 0.857±0.037 | 0.876±0.036 | 0.936±0.015 | 7.525±2.282 | 0.481±0.143 |
| SVC Classifier | 0.838±0.069 | 0.924±0.032 | 0.711±0.071 | 0.827±0.041 | 0.883±0.042 | 0.872±0.042 | 0.886±0.024 | 0.941±0.008 | **0.44±0.082** | 0.447±0.145 |
| XGBoost Classifier | **0.89±0.046** | **0.953±0.018** | 0.765±0.097 | 0.86±0.062 | 0.911±0.03 | 0.915±0.03 | **0.900±0.033** | 0.942±0.014 | 0.461±0.135 | 0.557±0.167 |
| XGBoost Random Forest Classifier | 0.857±0.064 | 0.936±0.029 | **0.773±0.057** | **0.868±0.04** | 0.885±0.047 | 0.907±0.039 | 0.891±0.041 | 0.936±0.011 | 1.79±0.853 | **0.558±0.105** |
| FedAvg LR | 0.69±0.16 | 0.874±0.042 | 0.617±0.109 | 0.772±0.054 | **0.955±0.037** | 0.818±0.054 | 0.863±0.026 | 0.935±0.008 | 0.655±0.14 | 0.278±0.25 |
| FedAvg MLP | 0.76±0.102 | 0.872±0.072 | 0.671±0.087 | 0.817±0.051 | 0.768±0.089 | 0.708±0.35 | 0.728±0.358 | 0.779±0.382 | 0.767±0.308 | 0.334±0.179 |
| FedAvg SGD | 0.828±0.048 | 0.92±0.025 | **0.757±0.048** | **0.904±0.049** | 0.707±0.033 | 0.871±0.032 | 0.872±0.018 | 0.939±0.008 | **0.545±0.032** | 0.47±0.084 |
| FedAvg XGBRF | **0.829±0.023** | **0.924±0.015** | 0.739±0.058 | 0.848±0.043 | 0.883±0.036 | **0.886±0.02** | 0.875±0.012 | 0.929±0.005 | 0.691±0.0 | **0.497±0.089w** |
| FedProx $\mu = 0.5$ LR | 0.755±0.142 | 0.887±0.041 | 0.653±0.088 | 0.791±0.042 | 0.941±0.031 | 0.704±0.349 | 0.729±0.358 | 0.784±0.384 | 0.609±0.155 | 0.362±0.198 |
| FedProx $\mu = 0.5$ MLP | 0.757±0.096 | 0.872±0.061 | 0.695±0.088 | 0.829±0.048 | 0.808±0.075 | 0.843±0.042 | 0.868±0.028 | 0.937±0.004 | 0.976±0.314 | 0.387±0.182 |
| FedProx $\mu = 2$ LR | 0.812±0.079 | 0.906±0.04 | 0.658±0.028 | 0.79±0.014 | 0.937±0.025 | 0.866±0.045 | **0.879±0.025** | **0.941±0.006** | 0.582±0.137 | 0.398±0.069 |
| FedProx $\mu = 2$ MLP | 0.765±0.079 | 0.868±0.06 | 0.694±0.069 | 0.83±0.042 | 0.798±0.045 | 0.706±0.348 | 0.724±0.355 | 0.781±0.382 | 0.9±0.368 | 0.379±0.133 |

**Table 2:** Performance of Several models trained using classical machine learning methods, and federated learning methods, where the number of participating clients in the federation is *N=2,* tested on the PDBP dataset. Data reported is mean and standard deviation across *K=6* fold cross validation.

| Algorithm Name | ROC-AUC | AUC-PR | Balanced Accuracy | Precision | Recall | F 0.5 | F 1 | F 2 | Log Loss | Matthews Correlation Coefficient |
| --- | --- | --- | --- | --- | --- | --- | --- | --- | --- | --- |
| AdaBoost Classifier | 0.834±0.021 | 0.891±0.015 | 0.697±0.026 | 0.757±0.023 | 0.917±0.033 | 0.835±0.016 | 0.834±0.009 | 0.905±0.004 | 0.639±0.005 | 0.456±0.029 |
| Bagging | 0.812±0.01 | 0.871±0.01 | 0.696±0.019 | 0.753±0.015 | 0.932±0.015 | 0.828±0.007 | 0.84±0.004 | 0.903±0.003 | 1.291±0.226 | 0.463±0.027 |
| GradientBoosting Classifier | 0.856±0.013 | 0.9±0.016 | 0.716±0.018 | 0.766±0.014 | 0.938±0.013 | 0.856±0.007 | **0.857±0.003** | **0.908±0.003** | **0.572±0.042** | 0.502±0.024 |
| KNeighbors Classifier | 0.586±0.024 | 0.735±0.019 | 0.551±0.019 | 0.664±0.011 | 0.946±0.024 | 0.707±0.008 | 0.783±0.001 | 0.899±0.0 | 3.322±0.641 | 0.169±0.042 |
| LinearDiscriminantAnalysis Classifier | 0.702±0.012 | 0.794±0.008 | 0.64±0.013 | 0.734±0.01 | 0.776±0.01 | 0.622±0.305 | 0.661±0.324 | 0.751±0.368 | 2.104±0.19 | 0.288±0.024 |
| LogisticRegression Classifier | 0.771±0.011 | 0.842±0.009 | 0.657±0.008 | 0.73±0.005 | 0.901±0.01 | 0.791±0.004 | 0.81±0.005 | 0.901±0.001 | 0.996±0.039 | 0.368±0.02 |
| MLP Classifier | 0.671±0.013 | 0.826±0.007 | 0.619±0.012 | 0.711±0.008 | 0.839±0.01 | 0.749±0.009 | 0.789±0.007 | 0.899±0.001 | 8.313±0.708 | 0.265±0.026 |
| QuadraticDiscriminantAnalysis Classifier | 0.525±0.022 | 0.721±0.022 | 0.525±0.022 | 0.671±0.024 | 0.366±0.097 | 0.688±0.0 | 0.779±0.0 | 0.898±0.0 | 18.716±1.33 | 0.05±0.042 |
| RandomForest | 0.736±0.006 | 0.825±0.005 | 0.524±0.005 | 0.649±0.003 | **0.985±0.005** | 0.764±0.007 | 0.792±0.005 | 0.899±0.0 | 0.596±0.004 | 0.132±0.025 |
| SGD Classifier | 0.662±0.017 | 0.845±0.007 | 0.65±0.016 | 0.728±0.011 | 0.878±0.024 | 0.758±0.01 | 0.803±0.007 | 0.898±0.0 | 10.11±0.525 | 0.343±0.034 |
| SVC Classifier | 0.701±0.007 | 0.808±0.004 | 0.593±0.011 | 0.693±0.006 | 0.844±0.02 | 0.742±0.004 | 0.793±0.002 | 0.901±0.001 | 0.65±0.019 | 0.214±0.029 |
| XGBoost Classifier | **0.862±0.008** | **0.905±0.007** | 0.719±0.021 | 0.77±0.016 | 0.932±0.013 | **0.864±0.006** | **0.857±0.003** | 0.906±0.003 | 0.691±0.031 | 0.504±0.03 |
| XGBoost Random Forest Classifier | 0.829±0.007 | 0.89±0.006 | **0.732±0.02** | **0.781±0.016** | 0.918±0.01 | 0.849±0.003 | 0.855±0.002 | 0.905±0.003 | 2.715±0.254 | **0.515±0.031** |
| FedAvg LR | 0.665±0.128 | 0.826±0.011 | 0.565±0.052 | 0.673±0.028 | **0.96±0.032** | 0.745±0.045 | 0.794±0.012 | 0.899±0.002 | 0.829±0.108 | 0.187±0.147 |
| FedAvg MLP | 0.69±0.018 | 0.78±0.012 | 0.629±0.01 | 0.719±0.007 | 0.828±0.022 | 0.744±0.008 | 0.791±0.007 | 0.899±0.001 | 1.038±0.239 | 0.282±0.024 |
| FedAvg SGD | 0.775±0.011 | 0.847±0.008 | 0.689±0.011 | **0.77±0.008** | 0.8±0.01 | 0.794±0.005 | 0.809±0.004 | 0.902±0.002 | **0.559±0.013** | 0.385±0.023 |
| FedAvg XGBRF | **0.794±0.007** | **0.876±0.009** | **0.695±0.023** | 0.754±0.017 | 0.919±0.012 | **0.825±0.007** | **0.838±0.008** | **0.902±0.003** | 0.691±0.0 | **0.451±0.035** |
| FedProx $\mu = 0.5$ LR | 0.704±0.101 | 0.823±0.015 | 0.584±0.042 | 0.683±0.022 | 0.943±0.03 | 0.762±0.038 | 0.795±0.009 | 0.9±0.002 | 0.866±0.092 | 0.232±0.115 |
| FedProx $\mu = 0.5$ MLP | 0.7±0.008 | 0.791±0.006 | 0.63±0.011 | 0.719±0.007 | 0.833±0.016 | 0.748±0.007 | 0.794±0.004 | 0.899±0.001 | 1.312±0.124 | 0.284±0.023 |
| FedProx $\mu = 2$ LR | 0.761±0.008 | 0.835±0.007 | 0.601±0.005 | 0.691±0.003 | 0.947±0.013 | 0.787±0.008 | 0.804±0.003 | 0.9±0.001 | 0.875±0.014 | 0.293±0.01 |
| FedProx $\mu = 2$ MLP | 0.695±0.022 | 0.785±0.015 | 0.631±0.02 | 0.722±0.013 | 0.818±0.02 | 0.747±0.014 | 0.791±0.005 | 0.899±0.001 | 1.231±0.285 | 0.282±0.044 |

### Sample dispersion among client sites negatively impacts global model performance

By splitting the training samples among an increasing number of clients, we aim to understand the implications of federation configurations that have more dispersed samples (**Fig. 4)**. In both the PPMI and PDBP data sets, there is a similar relative change in AUC-PR performance when increasing the number of client sites; the absolute performance scores, and variance are considerably higher for the PPMI test set than the PDBP test set. The performance of PPMI FedAvg XGBRF starts at 0.924 ± 0.015 AUC-PR in a federation of two client sites, and progressively drops to 0.861 ± 0.043 AUC-PR at 18 clients. For the PDBP performance, FedAvg XGBRF similarly reduces from 0.876 ± 0.009 to 0.752 ± 0.054 AUC-PR, at a minimum. Such a reduction in performance is also observed in the FedAvg SGD Classifier, which has an AUC-PR of 0.92 ± 0.025 for two clients, and an AUC-PR of 0.886 ± 0.055 for 18 clients in PPMI. In PDBP the same classifier starts at 0.847 ± 0.008 for two client sites, and ends at 0.798 ± 0.014 AUC-PR for 18 client sites. The trend of performance decline is observed for the LR classifiers as well, for both the PPMI and PDBP test sets. The FedAvg MLP, as well as both FedProx MLP classifiers do not exhibit such a reduction performance. In the PPMI test set, FedAvg MLP performance at *N=2* client sites is 0.872 ± 0.072 AUC-PR, and at *N=18* client sites, 0.876 ± 0.06 client sites. Similarly, for PDBP performance FedAvg MLP performance is 0.78 ± 0.012 for two client sites, and 0.781 ± 0.009 for *N=18* client sites. A nearly identical trend is observed in FedProx μ = 0.5, μ = 2 across both test sets. Detailed results are in supplementary materials, **Table 3**, and **Table 4**.

**Figure 4:**
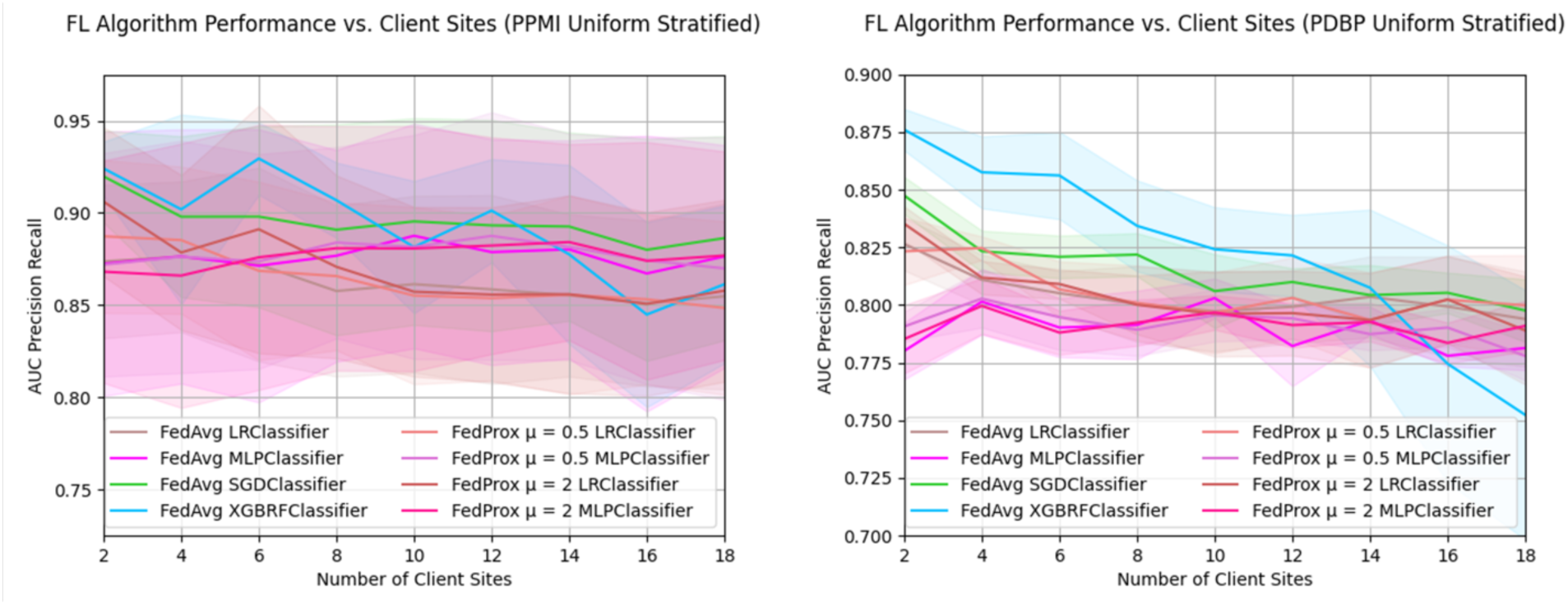
Sample dispersion among client sites negatively impacts global model performance. For a fixed training dataset, the AUC-PR of Federated Algorithms as the quantity of client sites increases. Training data is split uniformly among each member of the federation using stratified random sampling. The PDBP and PPMI datasets are used for external and internal validation, respectively. Presented data is mean score and standard deviation resulting from cross validation.

**Table 3:** AUC-PR score of models trained using Federated Learning as the quantity of client sites increased, tested on the PPMI dataset. Data reported is mean and standard deviation across *K=6* fold cross validation.

|  |  | Number of Clients |  |  |  |  |  |  |  |  |
| --- | --- | --- | --- | --- | --- | --- | --- | --- | --- | --- |
|  |  | 2 | 4 | 6 | 8 | 10 | 12 | 14 | 16 | 18 |
| Algorithm Name | FedAvg LR | 0.874 ± 0.042 | 0.876 ± 0.041 | 0.872 ± 0.053 | 0.858 ± 0.046 | 0.861 ± 0.048 | 0.859 ± 0.051 | 0.855 ± 0.044 | 0.851 ± 0.045 | 0.855 ± 0.05 |
| | FedAvg MLP | 0.872 ± 0.072 | 0.876 ± 0.069 | 0.871 ± 0.074 | 0.877 ± 0.057 | 0.888 ± 0.061 | 0.879 ± 0.061 | 0.88 ± 0.059 | 0.867 ± 0.075 | 0.876 ± 0.06 |
| | FedAvg SGD | 0.92 ± 0.025 | 0.898 ± 0.044 | 0.898 ± 0.049 | 0.891 ± 0.057 | **0.895 ± 0.056** | 0.893 ± 0.057 | **0.893 ± 0.051** | **0.88 ± 0.06** | **0.886 ± 0.055** |
| | FedAvg XGBRF | **0.924 ± 0.015** | **0.902 ± 0.051** | **0.929 ± 0.02** | **0.907 ± 0.02** | 0.882 ± 0.036 | **0.901 ± 0.028** | 0.878 ± 0.048 | 0.845 ± 0.05 | 0.861 ± 0.043 |
| | FedProx $\mu = 0$ LR | 0.887 ± 0.041 | 0.885 ± 0.04 | 0.869 ± 0.048 | 0.866 ± 0.04 | 0.855 ± 0.048 | 0.854 ± 0.045 | 0.856 ± 0.054 | 0.853 ± 0.046 | 0.849 ± 0.047 |
| | FedProx $\mu = 0$ MLP | 0.872 ± 0.061 | 0.876 ± 0.063 | 0.874 ± 0.058 | 0.884 ± 0.052 | 0.882 ± 0.061 | 0.888 ± 0.067 | 0.882 ± 0.061 | 0.874 ± 0.067 | 0.87 ± 0.071 |
| | FedProx $\mu = 2$ LR | 0.906 ± 0.04 | 0.879 ± 0.042 | 0.891 ± 0.067 | 0.871 ± 0.05 | 0.857 ± 0.046 | 0.856 ± 0.047 | 0.856 ± 0.054 | 0.851 ± 0.05 | 0.858 ± 0.049 |
| | FedProx $\mu = 2$ MLP | 0.868 ± 0.06 | 0.866 ± 0.072 | 0.876 ± 0.072 | 0.881 ± 0.066 | 0.881 ± 0.066 | 0.882 ± 0.059 | 0.884 ± 0.053 | 0.874 ± 0.064 | 0.877 ± 0.056 |

**Table 4:** AUC-PR score of models trained using Federated Learning as the quantity of client sites increased, tested on the PDBP dataset. Data reported is mean and standard deviation across *K=6* fold cross validation.

|  |  | Number of Clients |  |  |  |  |  |  |  |  |
| --- | --- | --- | --- | --- | --- | --- | --- | --- | --- | --- |
|  |  | 2 | 4 | 6 | 8 | 10 | 12 | 14 | 16 | 18 |
| Algorithm Name | FedAvg LR | 0.826 ± 0.011 | 0.811 ± 0.011 | 0.805 ± 0.01 | 0.8 ± 0.016 | 0.797 ± 0.017 | 0.799 ± 0.016 | 0.804 ± 0.018 | 0.799 ± 0.022 | 0.794 ± 0.021 |
| | FedAvg MLP | 0.78 ± 0.012 | 0.801 ± 0.014 | 0.79 ± 0.013 | 0.791 ± 0.015 | 0.803 ± 0.008 | 0.782 ± 0.017 | 0.793 ± 0.006 | 0.778 ± 0.005 | 0.781 ± 0.009 |
| | FedAvg SGD | 0.847 ± 0.008 | 0.823 ± 0.009 | 0.821 ± 0.009 | 0.822 ± 0.009 | 0.806 ± 0.016 | 0.81 ± 0.006 | 0.804 ± 0.013 | **0.805 ± 0.009** | 0.798 ± 0.014 |
| | FedAvg XGBRF | **0.876 ± 0.009** | **0.858 ± 0.016** | **0.856 ± 0.019** | **0.834 ± 0.02** | **0.824 ± 0.018** | **0.821 ± 0.018** | **0.807 ± 0.034** | 0.775 ± 0.051 | 0.752 ± 0.054 |
| | FedProx $\mu = 0$ LR | 0.823 ± 0.015 | 0.825 ± 0.005 | 0.807 ± 0.012 | 0.801 ± 0.016 | 0.797 ± 0.018 | 0.803 ± 0.018 | 0.793 ± 0.021 | 0.802 ± 0.019 | **0.8 ± 0.022** |
| | FedProx $\mu = 0$ MLP | 0.791 ± 0.006 | 0.803 ± 0.012 | 0.795 ± 0.014 | 0.789 ± 0.011 | 0.796 ± 0.011 | 0.794 ± 0.009 | 0.787 ± 0.007 | 0.79 ± 0.008 | 0.778 ± 0.011 |
| | FedProx $\mu = 2$ LR | 0.835 ± 0.007 | 0.812 ± 0.007 | 0.809 ± 0.006 | 0.8 ± 0.013 | 0.796 ± 0.018 | 0.796 ± 0.018 | 0.793 ± 0.02 | 0.802 ± 0.019 | 0.789 ± 0.023 |
| | FedProx $\mu = 2$ MLP | 0.785 ± 0.015 | 0.8 ± 0.012 | 0.788 ± 0.01 | 0.792 ± 0.01 | 0.797 ± 0.007 | 0.791 ± 0.01 | 0.793 ± 0.008 | 0.784 ± 0.009 | 0.791 ± 0.011 |

**Table 5:** The total runtime in seconds to train central and federated models, averaged over K folds. Algorithms are grouped by aggregation strategy (Central, FedAvg, FedProx). The lowest training time for each group is bolded.

| Algorithm Name | Runtime (Seconds) |
| --- | --- |
| LogisticRegression | 1.857e-02 ± 0.008 |
| SGDClassifier | **6.771e-03 ± 0.001** |
| MLPClassifier | 1.609e-01 ± 0.009 |
| XGBRFClassifier | 1.633e-01 ± 0.003 |
| FedAvg SGDClassifier | 1.513e+01 ± 1.497 |
| FedAvg XGBRFClassifier | 1.061e+01 ± 0.014 |
| FedAvg LRClassifier | **7.909e+00 ± 0.550** |
| FedAvg MLPClassifier | 8.755e+00 ± 0.141 |
| FedProx $\mu = 0.5$ LRClassifier | **8.747e+00 ± 0.158** |
| FedProx $\mu = 0.5$ MLPClassifier | 9.039e+00 ± 0.266 |
| FedProx $\mu = 2$ LRClassifier | 8.905e+00 ± 0.130 |
| FedProx $\mu = 2$ MLPClassifier | 9.260e+00 ± 0.163 |

### Data heterogeneity at client sites does not significantly influence model performance

To understand the implications of data heterogeneity among client sites, we examine the change in AUC-PR in a federation of two clients with respect to the split method (**Fig 5)**. In our experiments, we find that the performance changes introduced by dataset heterogeneity vary. Some FL models, such as FedAvg LR, FedProx μ = 0.5, FedAvg MLP, FedProx μ = 0.5 MLP, FedProx μ = 2 MLP exhibit performance improvements as both label heterogeneity and dataset size heterogeneity are introduced. The greatest performance improvement in PDBP is 0.018 AUCPR by FedProx μ = 2 MLP, and 0.029 by FedAvg LR in PPMI. Conversely, find that FedProx μ = 0.5 LR, FedProx μ = 2 LR, FedAvg SGD, FedAvg XGBRF exhibit performance degradation on account of dataset heterogeneity, where the greatest reduction in performance is 0.014 AUC-PR by XGBRF in PDBP, and 0.031 AUC-PR by FedProx μ = 2 LR on PPMI. In all such cases, the performance changes induced by dataset heterogeneity are marginal relative to other parameters such as algorithm choice or quantity of participating clients. The implications of performance heterogeneity as the number of clients increases are shown in **Supplementary Fig. 1, Supplementary Fig 2.**

**Figure 5:**
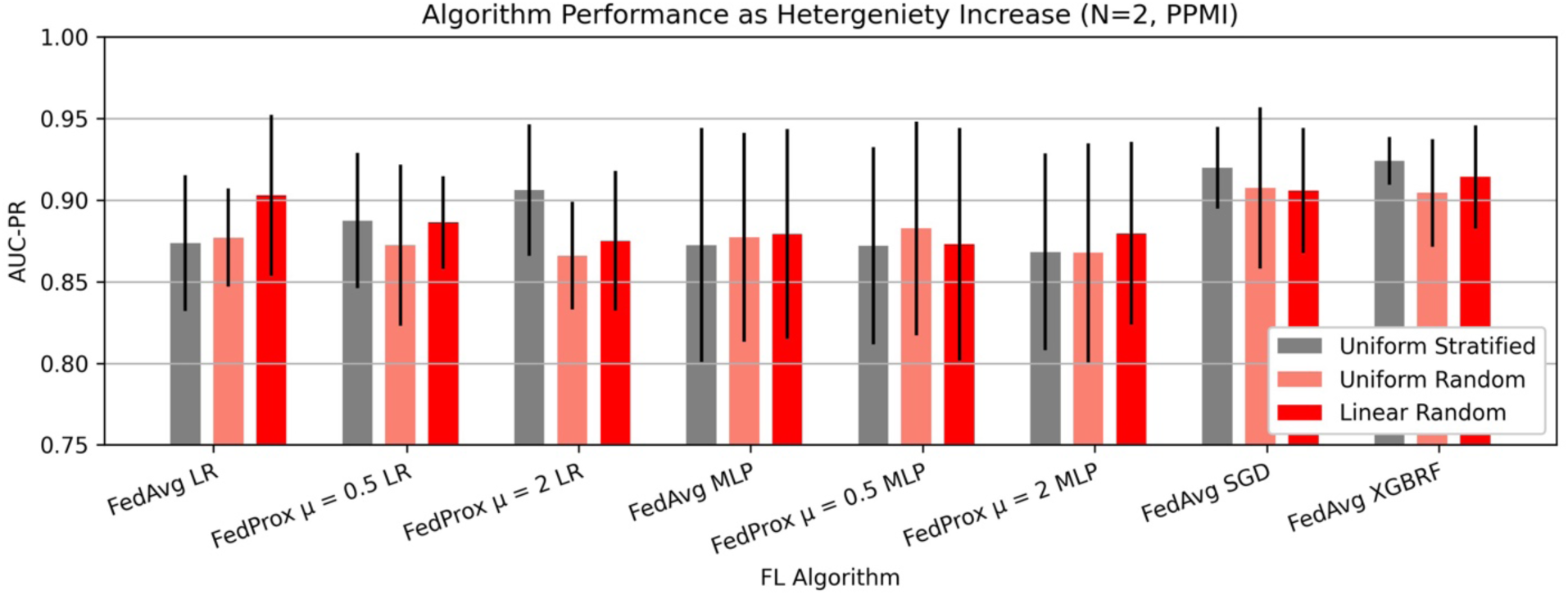
Data heterogeneity at client sites does not deeply influence model performance. The AUC-PR for a federation of 2 clients, for several split methods. Uniform stratified sampling, representing the most homogenous data distribution method, while uniform random, and linear random represent increasingly heterogeneous client distributions. Presented data is mean score and standard deviation resulting from cross validation.

### FL training time is not dramatically affected by choice in federated aggregation strategy

To shed light on the computational costs associated with using different FL aggregation strategies, we measure the model training time for FL algorithms in which the federation consists of N=2 client sites. For the FedAvg aggregation strategy, FedAvg LR had the lowest mean runtime of 7.909e+00 ± 0.550 seconds. Similarly, for the algorithms implementing FedProx aggregation, FedProx μ = 0.5 LRClassifier had the lowest overall runtime of 8.747e+00 ± 0.158 seconds. FedProx μ = 2 LRClassifier had the second lowest runtime for FedProx variants with a runtime of 8.905e+00 ± 0.130 seconds. For the MLP classifier, FedAvg, FedProx μ = 0.5, and FedProx μ = 2 had a progressively increasing runtimes 8.755e+00 ± 0.141 seconds, 9.039e+00 ± 0.266 seconds, 9.260e+00 ± 0.163 seconds respectively. FedAvg XGBRF, and FedAvg SGD had a considerably higher runtime of 1.061e+01 ± 0.014 and 1.513e+01 ± 1.497 seconds respectively. A visualization of the runtimes for FL algorithms is presented in **Figure 6**.

**Figure 6.**
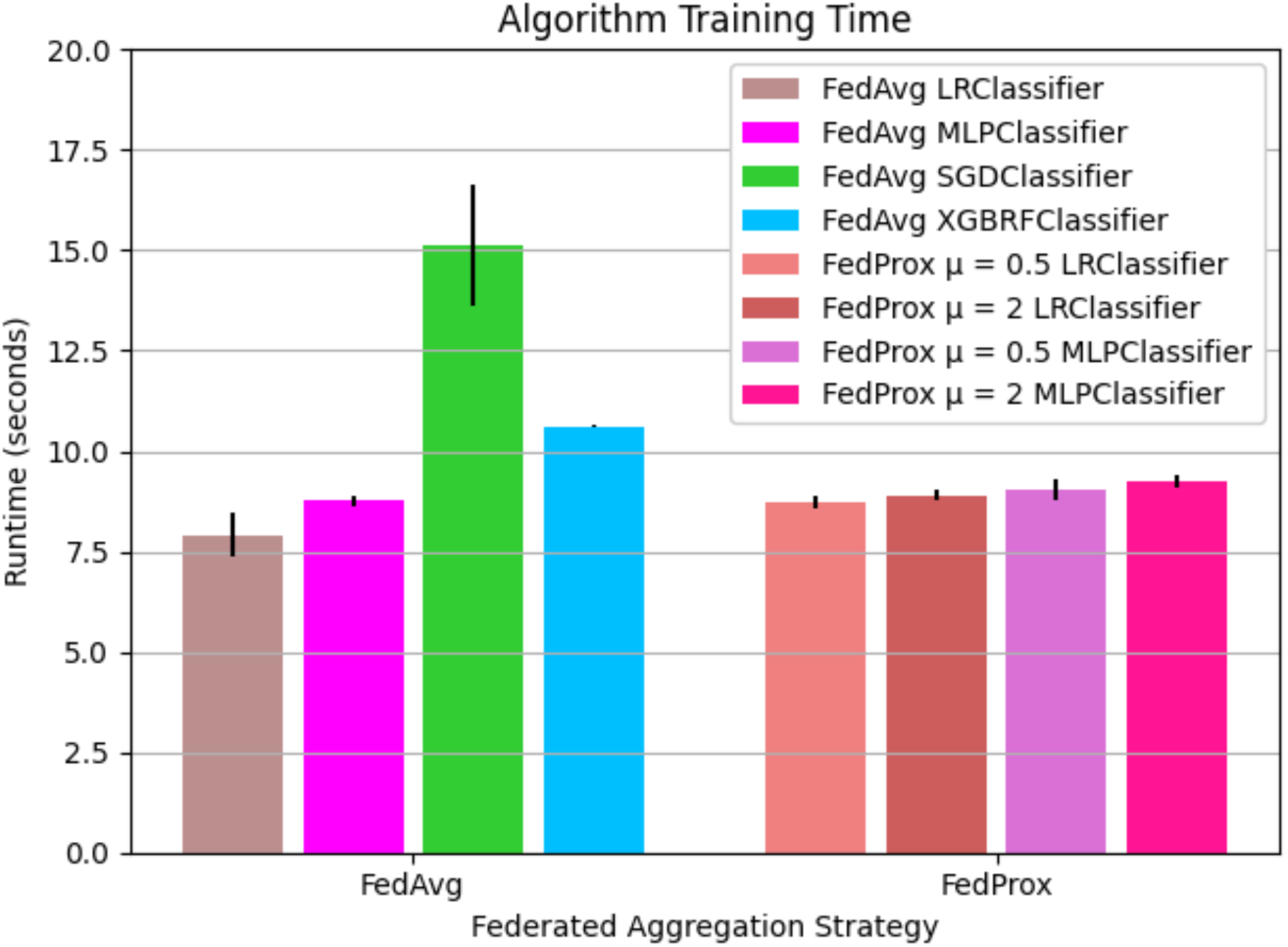
The mean runtime to train FL models using FedAvg and FedProx strategies. The mean total runtime in seconds to train FL models. FL models are trained on the PPMI training folds for 5 communication rounds. Algorithms are grouped by aggregation strategy. Results presented as mean and standard deviation over K=6 folds.

In contrast to the FL algorithms, central algorithm training time is at least an order of magnitude lower. The fastest model to train, SGD Classifier fits the model in 6.771e-03 ± 0.001 seconds, and the slowest model, MLP Classifier fits its model in 1.609e-01 ± 0.009. Central LR Classifier and XGBRF run in 1.857e-02 ± 0.008 and 1.633e-01 ± 0.003 seconds respectively. The full list of runtimes including central and federated algorithms is presented in **Table 7**.

## Discussion

In conjunction with increased access to genomic and transcriptomic data, the proliferation of high-quality Machine Learning open-source packages, has helped advance numerous long-standing challenges in biomedical research, such as disease subtyping, biomarker identification, and early disease diagnosis. The common bottleneck limiting such advances has thus shifted from the ability to apply ML methods to the availability of high-quality, well-designed datasets. Federated Learning has been cited as a promising means of alleviating the data scarcity problem through data-private collaborative model training ^24–26^. Previous works focus on applying FL to domains adjacent to multi-omics disease diagnosis, namely focusing on imaging data ^14, 27^, longitudinal health records ^15^, named entity recognition ^28^. Similar works such as ^29^ approach benchmarking FL models on biomedical datasets, but focus on comparing different FL aggregation strategies, rather than evaluating FL against a central baseline, as is done in our study. By approaching federated learning for multi-omics disease diagnosis from a performance benchmarking perspective, focusing on algorithms that are broadly accessible in the open-source community, we hope to shed light on what kind of practical performance can be achieved in a real-world setting where deep AI and Software Systems expertise may be limited. We additionally aim to understand what fundamental pitfalls researchers must be aware of before applying such methods in their multi-omics tasks.

When comparing centrally trained models against collaboratively trained models that implement the same local learner algorithm, our results indicate the FL trained model performance tends to be consistently less than that of the central method, while approximately following the performance of the central ML method. The general reduction in AUC-PR testing score between the FL and central method is noteworthy, but not a substantial deterioration. It can also be observed that for the studied aggregation strategies, FL model performance follows central model performance. In cases where the central model is performant, the FL trained model will be as well. In the case of the strongest central classifier in the central setting, XGBRF, the FL method implementing the same algorithm as a local learner, FedAvg XGBRF, also had the highest performance among models trained using FL. Additionally, we see that in many cases, FedAvg XGBRF outperforms central ML classifiers such as Logistic Regression, SGD, and MLP at the same task by a significant margin. This empirical result indicates that in cases where institutions must decide between applying FL methods to their setting, or centralizing data by complying with potentially stringent regulations, FL can be considered an effective option. In addition to this, because the implementation of such methods is available through open-source, strongly documented frameworks, the resource investment to achieve scientifically meaningful results may not be significant. We also note that because an FL model’s performance tracks, and seldom exceeds its central model performance, it can be crudely used to approximate the central model lower bound. Such an estimate of central performance, even if inexact, may be valuable for institutional stakeholders when deciding whether financial and administrative resources should be allocated to centralize several siloed datasets. We also note the overall reduction in performance between models trained using FL methods and models trained using central methods can be attributed to the federated aggregation process, which, in our case, is implemented as the unweighted average of the local learner model weights. Such a naive averaging process detracts from the parameter optimization implemented by local learners, but is a necessary cost to enable sample-private federated training. Furthermore, we note that several novel methods which implement more sophisticated weight aggregation strategies have been developed in academic settings, but are not always available as generally applicable open source packages. Overall, this test indicates that FL may be used to enable productive collaboration among institutions existing on opposite sides of geographic and policy boundaries, such as EU-GDPR, as well as across cloud providers and bare metal servers.

In the second arm of our study, we aim to understand the model performance cost of conducting collaborative training among a federation with increasing sample dispersion. Such a situation may arise, when institution stakeholders must comply with several layers of regulatory requirements, where centralizing some sites is easier than others. A concrete example of such a regulation is in the case of EU-GDPR, where transport of patient samples beyond the boundaries of the EU requires compliance with GDPR, and each country’s respective legislation mandates the transport of samples between countries within the EU. In this series of experiments, we assume that the globally available set of samples is constant, but the quantity of federation members containing the samples varies. Our experiments show that some methods, such as the LR classifiers, FedAvg SGD and FedAvg XGBRF, tend to exhibit performance degradation when there are more siloes with fewer samples per silo. We also observe that methods implementing MLP as a local learner, tend not to exhibit performance degradation with respect to sample concentration at siloes. Such methods do not necessarily achieve the best performance for any federation configurations, however, in the most extreme federation configuration of 18 clients, are still outperformed by methods such as FedAvg SGD and FedAvg LR. Methods whose performance is not strongly affected by silo size may represent practical starting points for the application of FL in an exploratory task. Ultimately, because FL models appear to have an optimal operating point which is modulated by the federation configuration, the final choice in FL methods used to reach peak performance should be determined by an exhaustive search. This finding suggests that a practical future format for applying FL in the biomedical setting may be through the auto-ML paradigm, which frameworks such as H2O ^30^, Auto Sklearn ^31^ currently implemented in the classical ML setting.

We additionally find in our studies that the implementation of heterogeneous client sites, with respect to dataset size and label counts, does not necessarily result in performance reductions for all algorithms. Some models such as FedAvg LR, models implementing MLP as a local learner, tend to increase performance, while models such FedAvg XGBRF, and FedAvg SGD exhibit performance degradation when the number of client sites is two (**Supplementary Figure 1**). We further find that when the number of clients is four, such heterogeneity has varying effects on performance, different from the configuration with two client sites. Overall, performance changes with respect to client dataset heterogeneity are marginal relative to changes introduced by factors such a number of clients per federation, or algorithm selection.

When comparing training time among federated learning algorithms, we found a mild progression in training time between FedAvg LRClassifier, FedProx μ = 0.5 LRClassifier, and FedProx μ = 2 LRClassifier respectively. The same trend can be observed for the FL algorithms using MLP as a local learner. The progressive increase in runtime may be attributed to the relative difference in complexity between the FedAvg and FedProx optimization mechanism. The objective of FedProx weight aggregation function includes a regularization term, μ, designed to handle heterogeneity among clients ^19^. The FedAvg optimization objective does not include this mechanism, making it conceptually simpler. In the context of this study, the performance differences incurred by choice in aggregation method are minor relative to parameters such as choice in local learner, number of federated aggregation rounds, and dataset size.

When comparing the training time of central and federated models, we find that the central model training time is at least an order of magnitude lower than the federated training time. This result is not surprising, given that federated models implement global weight aggregation and updating steps in model training. Given that our study performs federated optimization in simulation, production deployments of FL methods can be expected to have slower overall runtimes due to network latency and operating system throughput capabilities.

The algorithms used in evaluating collaboratively trained models using FL against centralized applications of their local learner methods are detailed in **Supplementary Table 2**. In our study, we omitted using closed-source FL methods available through platform interfaces since these methods allow data governance capabilities to external parties, or vendor security evaluations, which in some cases instantiates barriers to productive research. While numerous publications explore methodological improvements that push forward state-of-the-art FL model performance in an experimental setting, we encountered challenges in applying such methods in our case, as many of these academic studies do not result in broadly applicable packages. In our research, we found a set of open-source projects that implement FL methods and provide out-of-the-box solutions, or well-designed examples that could be interpolated to the multi-omics classification task to be limited. Ultimately, the FL interfaces made available by NVFlare ^32^ and Flower ^33^ were selected to conduct experiments, with local learners implemented using Sklearn ^22^ and DMLC ^21^ packages. Several open-source projects, such as Owkin ^34^, Tensorflow Federated ^35^, and OpenFL ^36^ provide full interfaces for implementing deep learning models in TensorFlow ^35^, and PyTorch ^37^, but such deep methods are less suitable for tabular tasks on datasets with only a handful of samples, as is the case the multi-omics datasets used in this study. Additionally, we found that while several packages provided abstract interfaces for implementing any arbitrary set of local learners and aggregation strategies, without detailed examples with a straightforward path to adaptation to a particular research task, the practical application of such methods becomes challenging, and less approachable for groups which may be resource constrained.

The extent to which Federation Site configurations could be studied was largely limited by the number of case patients within the dataset. Concretely, the implications of heterogeneity in site data could only be observed to the extent that each silo would maintain enough samples from case and control cohorts to allow the local learner to successfully train. Datasets at silos needed to have at least one sample from both the case and control groups. Similarly, although the PPMI dataset was collected across several geographically distributed institutions, point of origin information is not available for each sample, preventing the evaluation of performance on naturally occurring siloes. In our study, all experiments assume that the collective dataset available among all client sites has a constant size. An additional limitation of our work is in observing the effect of adding federation members, which contribute novel samples to the federation.

While FL methods enable data owners to maintain governance of their local datasets, on its own, FL does not provide end to end privacy guarantees. Our study examines the utility of FL methods in the multi-omics case study to understand the availability and characteristics of FL, and does not include a concrete evaluation of privacy, or security methods. Thus, we assume that the federated learning aggregation server is neither dishonest, curious, nor malicious in any way and fulfills its functions as an intermediary between client sites benevolently. Privacy preserving methods orthogonal to FL such as Differential Privacy (DP) enable the application of FL with formal guarantees of sample-privacy ^38^. Such approaches were not included in the scope of this evaluation, but represent a factor which should be considered when applying FL methods in settings where verifiable sample privacy guarantees are critical. In our experimentation, we do not focus on the implications of the federation which has heterogeneous compute capabilities, since applying machine learning model fitting on datasets with few samples can be done without much difficulty.

The datasets utilized in our analysis, including PPMI and PDBP, are sourced from the Accelerating Medicines Partnership Parkinson’s Disease (AMP-PD) initiative. This initiative plays a pivotal role in unifying transcriptomic and genomic samples, ensuring consistency and accuracy through central harmonization and joint-calling processes. Furthermore, the construction of machine learning features for our analysis is also centralized, leveraging these cohesive datasets. Recognizing the potential for broader application, our future focus includes exploring federated analysis tasks ^39–41^. This involves enhancing cross-silo harmonization, joint-calling, and feature construction across diverse datasets. To facilitate this, the development of specialized federated learning libraries, specifically tailored for genomics and transcriptomics, is crucial. Such advancements will not only democratize access to federated learning (FL) methods for the wider biomedical community but also significantly broaden the scope for applying machine learning techniques in various biomedical contexts.

Overall, we believe that this work sheds light onto the feasibility, and noteworthy characteristics of applying Federated Learning for omics analysis. Through our experiments, we find that collaboratively trained FL models can achieve high classification accuracy in multi-omics Parkinson’s Disease Diagnosis, and can remain relatively performance despite heterogeneity among client sites. We also find in our evaluation that although FL is a relatively novel research space in bioinformatics, there is sufficient access to open source methods which biomedical researchers may leverage to enable productive collaborations.

## Experimental Procedures

### Resource Availability

#### Lead Contact

Further information and requests for resources and reagents should be directed to and will be fulfilled by the lead contact, Faraz Faghri.

#### Materials Availability

This study did not generate new unique materials.

#### Data and Code Availability

##### Data

The data used in this study was access-controlled from the Parkinson’s Progression Marker Initiative (PPMI, http://www.ppmi-info.org/) and the Parkinson’s Disease Biomarkers Program (PDBP, https://pdbp.ninds.nih.gov/).

##### Code

To facilitate replication and expansion of our work, we have made the notebook publicly available in an open repository^42^. It includes all code, figures, models, and supplements for this study. The code is part of the supplemental information; it includes the rendered Jupyter notebook with full step-by-step data preprocessing, statistical, and machine learning analysis.

Any additional information required to reanalyze the data reported in this paper is available from the lead contact upon request.

#### Datasets

The dataset used in this study as the basis for training and as the internal test set is the Parkinson’s Progression Marker Initiative (PPMI) dataset. The PPMI dataset represents a longitudinal, observational study where patients contribute clinical, demographic, imaging data, biological samples for whole-genome sequencing, and whole-blood RNA sequencing. Samples are collected at 33 clinical sites globally and across a time span of anywhere from 5 to 13 years. This preprocessed dataset consists of 171 samples of case patients diagnosed with Parkinson’s disease and 427 healthy control patients. The PPMI cohort contains newly diagnosed, and drug-naive patient samples. The PPMI cohort contains 209 (36%) female samples, 388 (67%) male samples (**Supplementary Table 4**).

The dataset used in this study for external, out-of-distribution validation is the Parkinson’s Disease Biomarker Program (PDBP). It is a longitudinal, observational study where patients contribute clinical, demographic, and imaging data and biological samples for whole-genome and whole-blood RNA sequencing. The preprocessed dataset consists of 712 healthy control patients and 404 case patients diagnosed with Parkinson’s disease. Each sample comprises 713 features, including genetic, transcriptomic, and clinico-demographic information collected at the baseline. The PDBP cohort consists of 480 (43%) female samples, and 636 (57%) male samples. (**Supplementary Table 5**).

Both PPMI and PDBP data used in this study were acquired through the AMP-PD initiative ^43^, an effort to provide harmonized datasets that include common clinical and genomic data. Through this initiative, the PPMI and PDBP datasets are centrally joint-called and harmonized to allow standardization across cohorts.

Transcriptomic data from whole blood RNA sequencing was generated by the Translational Genomics Research Institute team using standard protocols for the Illumina NovaSeq technology and processed through variance-stabilization and limma pipelines ^44^ for experimental covariates. Gene expression counts for protein-coding genes were extracted, then differential expression *p* values were calculated between cases and controls using logistic regression adjusted for additional covariates of sex, plate, age, ten principal components, and percentage usable bases. A comprehensive description of the RNA-Sequencing method is presented in ^45^ for PPMI, and ^46^ for PDBP.

For genetic data, sequencing data were generated using Illumina’s standard short-read technology, and the functional equivalence pipeline during alignment was the Broad Institute’s implementation ^47^. Applied quality control measures included criteria like gender concordance and call rate, with a focus on SNPs meeting the GATK gold standards pipeline and additional filters like non-palindromic alleles and missingness by case-control status thresholds. Polygenic risk scores (PRS) were constructed using effect sizes from a large European genome-wide meta-analysis, supplementing the genetic data from whole genome sequences. The process from sample prep to variant calling is comprehensively described in ^43^.

Quality control for genetic samples based on genetic data output by the pipeline included the following inclusion criteria: concordance between genetic and clinically ascertained genders, call rate > 95% at both the sample and variant levels, heterozygosity rate < 15%, free mix estimated contamination rate < 3%, transition:transversion ratio > 2, unrelated to any other sample at a level of the first cousin or closer (identity by descent < 12.5%), and genetically ascertained European ancestry. For inclusion of whole-genome DNA sequencing data, the variants must have passed basic quality control as part of the initial sequencing effort (PASS flag from the joint genotyping pipeline) as well as meeting the following criteria: non-palindromic alleles, missingness by case-control status *P* > 1E-4, missingness by haplotype *P* > 1E-4, Hardy– Weinberg *p* value > 1E-4, minor allele frequency in cases > 5% (in the latest Parkinson’s disease meta-GWAS by ^48^). As an a priori genetic feature to be included in our modeling efforts, we also used the basic polygenic risk score from the latest Parkinson’s disease meta-GWAS (genome-wide significant loci only) that did not include our testing or training samples as weights ^48^.

Compared to the PPMI dataset, PDBP includes an additional 40 genetic features, which are excluded from this study, allowing PPMI and PBDP to have the same feature set. Additionally, the PPMI samples are collected before any medical intervention, whereas the PDBP samples are, in some cases, collected after patient treatment has commenced. Since the PDBP dataset may include artifacts which result from disease treatment, the PDBP dataset is used exclusively for evaluation to avoid the possibility of label leakage. A shortened version of the final feature set is provided in **Supplementary Table 1**. A comprehensive feature list is available in the external code repository ^42^.

Each sample consists of 673 features, including genetic, transcriptomic, and clinico-demographic information collected at the baseline. Of the 673 features, 72 originate from genome sequencing data and polygenic risk score, 596 are transcriptomic, and 5 are clinico-demographic. The clinico-demographic features include age, family history, inferred Ashkenazi Jewish status, sex, University of Pennsylvania Smell Identification (UPSIT) score.

#### Data Preprocessing

The construction of features from genomic, transcriptomic, and clinico-demographic data is handled for each cohort independently, and in a centralized manner, for the entirety of the cohort. As part of the initial data preprocessing, principal components summarizing genetic variation in DNA and RNA sequencing data modalities are generated separately. For the DNA sequencing, ten principal components were calculated based on a random set of 10,000 variants sampled after linkage disequilibrium pruning that kept only variants with r^2^ < 0.1 with any other variants in ±1 MB. As a note, these variants were not p value filtered based on recent GWAS, but they do exclude regions containing large tracts of linkage disequilibrium ^49^. Our genetic data pruning removed SNPs in long tracts of high LD such as in the HLA region (we excluded any SNPs within r^2^ > 0.1 within a sliding window of 1 MB), while retaining known genetic risk SNPs within the region. For RNA sequencing data, all protein-coding genes’ read counts per sample were used to generate a second set of ten principal components. All potential features representing genetic variants (in the form of minor allele dosages) from sequencing were then adjusted for the DNA sequence-derived principal components using linear regression, extracting the residual variation. This adjustment removes the effects of quantifiable European population substructure from the genetic features prior to training, this is similar in theory to adjusting analyses for the same principal components in the common variant regression paradigm employed by GWAS models. The same was done for RNA sequencing data using RNA sequencing derived principal components. This way, we statistically account for latent population substructure and experimental covariates at the feature level to increase generalizability across heterogeneous datasets. In its simplest terms, all transcriptomic data were corrected for possible confounders, and the same is done for genotype dosages. After adjustment, all continuous features were then Z-transformed to have a mean of 0 and a standard deviation of 1 to keep all features on the same numeric scale when possible. Once feature adjustment and normalization were complete, internal feature selection was carried out in the PPMI training dataset using decision trees (extraTrees Classifier) to identify features contributing information content to the model while reducing the potential for overfitting prior to model generation ^22^ ^50^. Overfitting here is defined as the over-performance of a model in the training phase with minimal generalizability in the validation dataset due to the inclusion of potentially correlated or unimportant features. The implementation of decision trees for feature selection helps remove redundant and low-impact features, helping us to generate the most parsimonious feature set for modeling. Feature selection was run on combined data modalities to remove potentially redundant feature contributions that could artificially inflate model accuracy. Export estimates for features most likely to contribute to the final model in order of importance were generated by the extraTrees classifier for each of the combined models, and are available on the Online Repository. By removing redundant features, the potential for overfitting is limited while also making the models more conservative. Additionally, if a variant provided redundant model information, such as being in strong linkage with a PRS variant, it would be removed from the potential feature list.

Feature selection was performed using the extremely randomized trees classifier algorithm, extraTrees ^50^, on combined data modalities to remove redundant feature contributions that could overfit the model to optimize the information content from the features and limit artificial inflation in predictive accuracy that might be introduced by including such a large number of features before filtering. In many cases, including more data might not be better for performance. With this in mind, we attempted to build the most parsimonious model possible using systematic feature selection criteria ^51^. Among the top 5% of features ranked in the Shapley analysis, the mean correlation between features was r2 < 5%, with a maximum of 36%. By removing redundant features using correlation-based pruning and an extraTrees classifier as a data munging step, the potential for overfitting is limited while also making the models more conservative.

Clinical and demographic data ascertained as part of this project included age at diagnosis for cases and age at baseline visit for controls. Family history (self-reporting if a first or second-degree relative has a PD diagnosis) was also a known clinico-demographic feature of interest. Ashkenazi Jewish status was inferred using principal component analysis comparing those samples to a genetic reference series, referencing the genotyping array data from GSE23636 at Gene Expression Omnibus as previously described elsewhere ^43, 52^. Sex was clinically ascertained but also confirmed using X chromosome heterozygosity rates. The University of Pennsylvania smell inventory test (UPSIT) was used in modeling ^53^. A comprehensive description of data and preprocessing is described in ^6^.

#### Quantification and Statistical Analysis

We conducted K-fold cross-validation on the PPMI dataset, where *K=6*, allowing each fold to contain approximately 100 samples. For each cross-validation fold, *1/K* of the PPMI samples are withheld as a holdout test set. The remaining training split of *K - 1/K* samples are further split using uniform stratified random sampling at an 80:20 ratio into training and validation subsets. The evaluation set was used for cross validated hyperparameter tuning in the central and federated models. The PPMI dataset was selected as the training and internal test set due to the fact that patients samples recorded in the PPMI protocol are newly diagnosed and drug-naive. Additionally, by training on the PPMI dataset, the model is developed on patients earlier on in their disease course. This intentional choice was made in the hopes the model would identify other individuals early on in their disease course and prioritize them for follow-up. The PDBP cohort samples are collected within five years after diagnosis, and may be actively taking medications. While the PDBP cohort is larger, because samples are collected several years after diagnosis, and because patients may be actively taking medication, there is a possibility of label leakage, ultimately motivating the usage of the PPMI dataset for training. Due to the similar nature of the PPMI and PDBP dataset, after processing, the PDBP dataset can be used as an external test set, approximating out-of-distribution model performance.

To conduct federated model training, the fully preprocessed PPMI dataset is split into disjoint client subsets, using one of the split strategies, and assigned to a local learner. To train a global model using the data among all federation participants, an iterative optimization process is run for a predefined set of rounds (**Fig. 2**). During this process, federation members fit local learner models to their locally available datasets. The parameters resulting from local model fitting are then sent to the central aggregation server. Once all model parameters are received, the aggregation server applies a federated learning strategy to the set of model weights, resulting in aggregated model weights, referred to as the global model. The global model is then sent back to the client sites, and used as the starting point for local learner optimization in subsequent iterations. The best performing global model on the evaluation set is used for final testing.

We simulate two types of heterogeneity in our experiments by distributing samples from the training dataset using different split methods. The split method uniform stratified sampling implements label, and site size homogeneity. The split method uniform random sampling implements label heterogeneity, but site size homogeneity. The split method linear random sampling implements label and site size heterogeneity (**Fig. 3**).

Local client data sets were formed by applying such sampling techniques to the centralized training data set. Uniform stratified sampling was used to form the *K* folds and during experiments that study homogenous federations. Uniform stratified sampling entails sampling the source dataset such that there is a nearly even distribution of samples among each of the *N* clients, and the ratio of cases to controls across each client subset is equivalent (where surplus samples are assigned to one of the client sites at random). This method was implemented by partitioning the source dataset by phenotype and then, without replacement, assigning *1/N* samples of each phenotype partition to each client using uniform random sampling, with the last client receiving any extra samples. In practice, this additional data was less than 10 of samples. Uniform random sampling entails assigning *1/N* using uniform random sampling. Linear random sampling entails assigning *c_i_* samples to a client site, where the following is true:

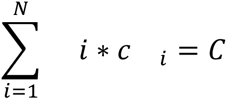

In the above formula, C is the number of samples to distribute, in practice the size of the PPMI training set, and *i* is the index of the client site. As with previous methods, the final client receives any surplus samples left over. The effect of this linear random sampling strategy is that each of the *N* clients contains an increasing number of samples relative to the previous clients, and each client site contains a random distribution of cases versus controls.

To measure algorithm runtime, for central algorithms, we measured the quantity of seconds from model initialization, to model training completion. For federated algorithms, we measured the quantity of seconds from model initialization, to the end of the FL training simulation. For FL models, model optimization was conducted for a federation of *N=2* federation clients, for 5 aggregation rounds.

In our simulation configurations, federation rounds operate synchronously, and without failure. Hyperparameters that were used to compute the final results are reported in the appendix, including the random seed used for the presented results.

The Federated Machine Learning methods implemented in the study utilize the federated aggregation methods FedAvg ^18^, FedProx ^19^, and the local learner classification methods, Logistic Regression ^17^, Multi-Layer Perceptron ^20^, Stochastic Gradient Descent ^23^, and XGBoost ^21^ available through Sklearn ^22^ and DMLC ^21, 22^. The aggregation methods are implemented using NVFlare ^32^, and Flower ^33^, while local learner methods are implemented using Sklearn, and DMLC APIs. Configuration details are available in the supplementary section. Simulation Frameworks used to implement model experiments are made available through the NVFlare and Flower packages. A single client site exhibits a minor computational cost of a single GB CPU and a single logical processor, which must be available throughout the life of the simulation. The simulations for both NVFlare and Flower required 18 Gb of RAM, and 18 logical cores. A simulation to train a single FL model takes less than 1 minute to complete. Running the full suite of simulations to reproduce the paper figures takes 6-8 hours. All experiments were conducted on Redhat Enterprise Linux Distribution.

## Acknowledgments

We thank the patients and their families who contributed to this research. This research was supported in part by the Intramural Research Program of the National Institute on Aging (NIA) and National Institute of Neurological Disorders and Stroke (NINDS), both part of the National Institutes of Health, within the Department of Health and Human Services; project number ZIAAG000534 and the Michael J Fox Foundation. Data used in the preparation of this article were obtained from the AMP PD Knowledge Platform. For up-to-date information on the study, please visit https://www.amp-pd.org. AMP PD—a public-private partnership—is managed by the FNIH and funded by Celgene, GSK, the Michael J. Fox Foundation for Parkinson’s Research, the National Institute of Neurological Disorders and Stroke, Pfizer, Sanofi, and Verily. Clinical data and biosamples used in the preparation of this article were obtained from the Parkinson’s Progression Markers Initiative (PPMI), and the Parkinson’s Disease Biomarkers Program (PDBP). PPMI—a public-private partnership—is funded by the Michael J. Fox Foundation for Parkinson’s Research and funding partners, including full names of all of the PPMI funding partners found at http://www.ppmi-info.org/fundingpartners. The PPMI Investigators have not participated in reviewing the data analysis or content of the manuscript. For up-to-date information on the study, visit http://www.ppmi-info.org. The Parkinson’s Disease Biomarker Program (PDBP) consortium is supported by the National Institute of Neurological Disorders and Stroke (NINDS) at the National Institutes of Health. A full list of PDBP investigators can be found at https://pdbp.ninds.nih.gov/policy. The PDBP Investigators have not participated in reviewing the data analysis or content of the manuscript. PDBP sample and clinical data collection is supported under grants by NINDS: U01NS082134, U01NS082157, U01NS082151, U01NS082137, U01NS082148, and U01NS082133.

## Author Contributions

B.D., M.B.M, A.D., D.V., M.A.N., J.S., and F.F. contributed to the concept and design of the study. B.D., M.B.M, A.D., D.V., M.A.N., and F.F were involved in the acquisition of data, data generation, and data cleaning. B.D., A.D., M.A.N., and F.F. did the analysis and interpretation of data. B.D., M.B.M, A.D., D.V., P.S.L, M.A.N., J.S., and F.F. contributed to the drafting of the article and revising it critically.

## Inclusion and Diversity Statement

The research team is comprised of diverse racial, national, and gender identity groups. Study has incorporated clinical and genomic data from the PPMI and PDBP projects as well as including its diverse group of investigators across career stages to collaborate on this project.

