## Supplemental Materials for "Federated Learning for multi-omics: a performance evaluation in Parkinson’s disease"

### Supplementary Information

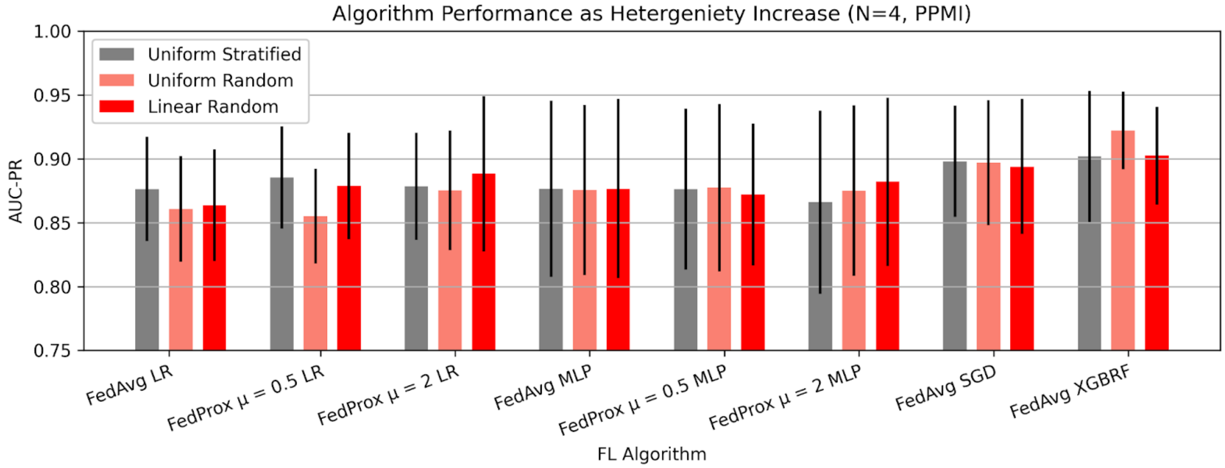


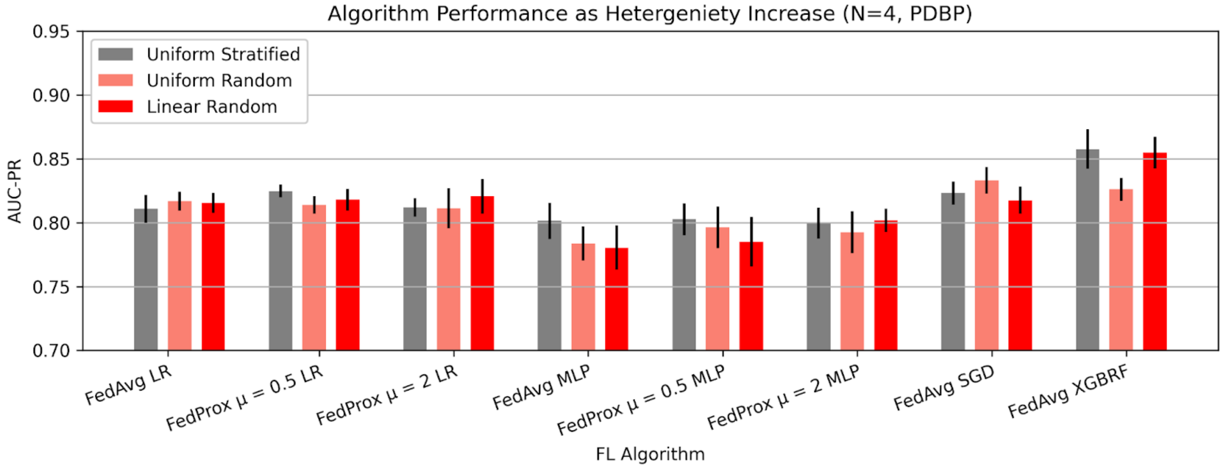


**Supplementary Figure 1: The AUC-PR for a federation of 4 clients, for several split methods.**

Uniform stratified sampling, representing the most homogenous data distribution method, while uniform random, and linear random represent increasingly heterogeneous client distributions. Presented data is mean score and standard deviation resulting from cross validation.

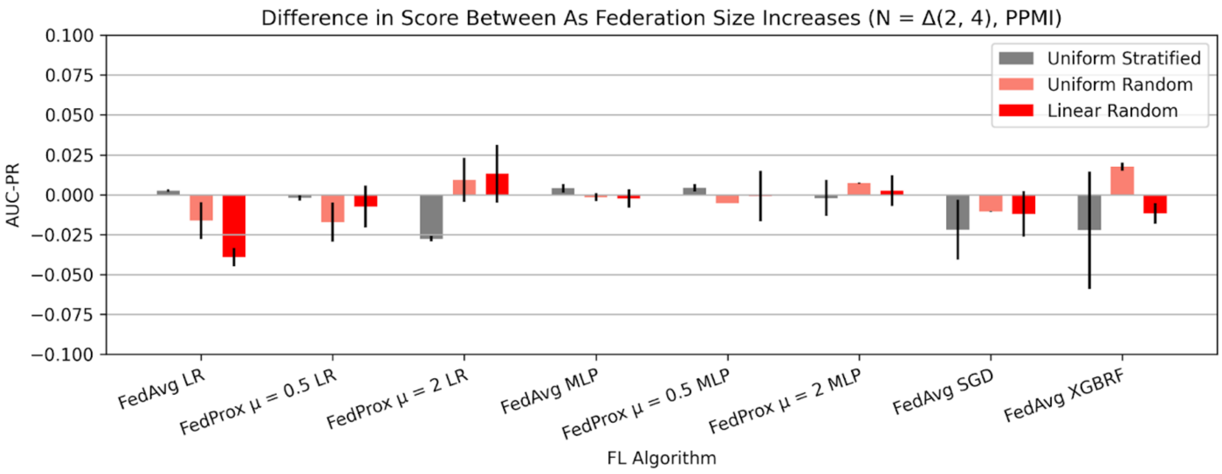


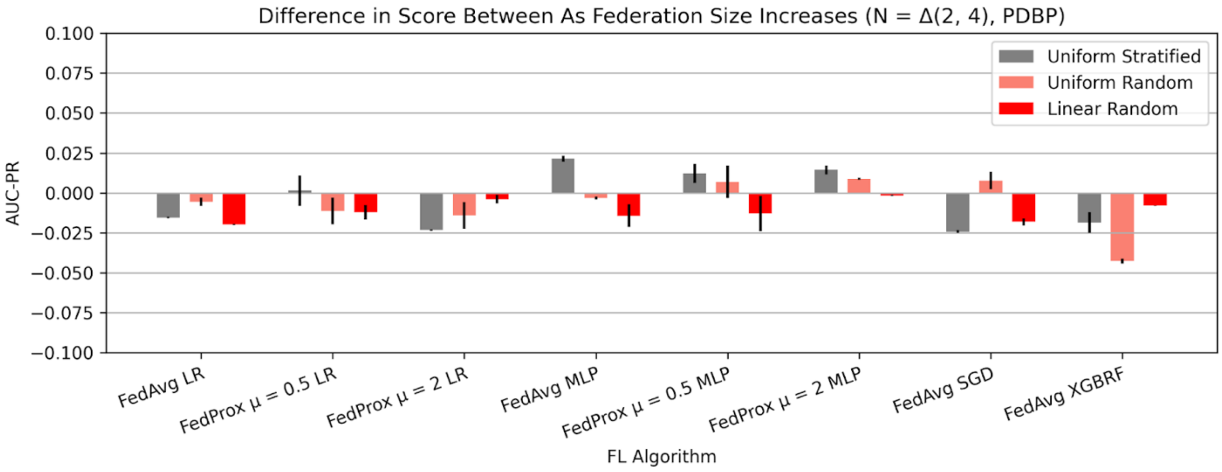


**Supplementary Figure 2: The difference between AUC-PR scores between 2 and 4 clients, for several split methods.**

Uniform stratified sampling, representing the most homogenous data distribution method, while uniform random, and linear random represent increasingly heterogeneous client distributions. Presented data is mean score and standard deviation resulting from cross validation.

| Feature Name | Feature Source |
|---|---|
| Age | Clinico-demographic |
| Family History | Clinico-demographic |
| Male | Clinico-demographic |
| UPSIT | Clinico-demographic |
| Inferred Ashkenazi Jewish | Clinico-demographic |
| PRS90 | Genetic |
| rs10182170 | Genetic |
| rs10186643 | Genetic |
| ENSG00000000938 | Transcriptomic |
| ENSG00000001629 | Transcriptomic |
| ENSG00000008394 | Transcriptomic |

**Supplementary Table 1:** The name and source of all clinic-demographic features, the first three genetic features, and the first three transcriptomic features. The comprehensive list of 674 features is available in the supplementary code repository.

| Algorithm Name | Central Learner API | Federated Local Learner API | Federated Weight Aggregation Method | Federated Learning API |
|---|---|---|---|---|
| Logistic Regression | Scikit Learn | Scikit Learn | FedAvg, FedProx | Flower Framework |
| MLP Classifier | Scikit Learn | Scikit Learn | FedAvg, FedProx | Flower Framework |
| SGD | Scikit Learn | Scikit Learn | FedAvg | NVIDIA Flare |
| RF XGBoost Classifier | DMLC | DMLC | FedAvg | NVIDIA Flare |

**Supplementary Table 2:** The description of frameworks used to implement central, and federated learning models.

| | FedAvg LRClassifier | FedAvg MLPClassifier | FedAvg SGDClassifier | FedAvg XGBRFClassifier | FedProx μ = 0 LRClassifier | FedProx μ = 2 LRClassifier | FedProx μ = 0 MLPClassifier | FedProx μ = 2 MLPClassifier |
|---|---|---|---|---|---|---|---|---|
| **LogisticRegression** | greater* | greater* | greater | lesser | greater* | greater* | greater* | greater* |
| **MLPClassifier** | lesser* | lesser | lesser* | lesser* | lesser* | lesser* | lesser | lesser* |
| **SGDClassifier** | lesser* | lesser | lesser* | lesser* | lesser* | lesser* | lesser | lesser* |
| **XGBRFClassifier** | greater* | greater* | greater* | greater* | greater* | greater* | greater* | greater* |

**Supplementary Table 3:** Method comparison table indicating statistical significance of the observed differences (greater, lesser) in performance measure (ROC-AUC) between models fit using central and federated methods on the external test set. Significance determined using DeLong's test, where an asterisk indicates statistical significance ($p < 0.05$).

| | Case | Control | Total |
|---|---|---|---|
| Male | 109 | 279 | 388 |
| Female | 62 | 147 | 209 |
| Total | 171 | 426 | 597 |

**Supplementary Table 4:** Value counts of PPMI cohort.

| | Case | Control | Total |
|---|---|---|---|
| Male | 185 | 451 | 636 |
| Female | 219 | 261 | 480 |
| Total | 404 | 712 | 1116 |

**Supplementary Table 5:** Value counts of PDBP cohort.

| Class Name | Algorithm Package | Package Version |
|---|---|---|
| LogisticRegression | sklearn | 1.3.0 |
| RandomForestClassifier | sklearn | 1.3.0 |
| AdaBoostClassifier | sklearn | 1.3.0 |
| GradentBoostingClassifier | sklearn | 1.3.0 |
| SGD | sklearn | 1.3.0 |
| SVC | sklearn | 1.3.0 |
| MLPClassifier | sklearn | 1.3.0 |
| KNNClassifier | sklearn | 1.3.0 |
| LinearDiscriminantAnalysis | sklearn | 1.3.0 |
| BaggingClassifier | sklearn | 1.3.0 |
| XGBClassifier | xgboost | 1.7.6 |
| XGBRFClassifier | xgboost | 1.7.6 |

**Supplementary Table 6:** The class names, algorithm packages, and package versions used to implement local learners in federated models, and central machine learning models.
